# Unconscious learning and automatic inhibition are accompanied by frontal theta and sensorimotor interactions

**DOI:** 10.1101/777961

**Authors:** Silvia L. Isabella, J. Allan Cheyne, Douglas Cheyne

## Abstract

Cognitive control of behavior is often accompanied by theta-band activity in the frontal cortex, and is crucial for overriding habits and producing desired actions. However, the functional role of theta activity in controlled behavior remains to be determined. Here, we used a behavioral task (Isabella et al., 2019) that covertly manipulated the ability to inhibit (and switch) motor responses using a repeating pattern of stimuli that reduced reaction times (RT) to probable over unexpected stimuli, without participants’ awareness of the pattern. We combined this task with concurrent measures of brain activity and pupil diameter (as a measure of cognitive activity) of 16 healthy adults during response preparation and inhibition during changes in stimulus probability. Observed RT provided evidence of pattern learning and pupillometry revealed parametric changes in cognitive activity with stimulus probability. Critically, reliable pupillary effects (Hedge’s *g* = 1.38) in the absence of RT differences (*g* = 0.10) indicated that cognitive activity increased without overt changes in behavior (RT). Such increased cognitive activity was accompanied by parametric increases in frontal theta and sensorimotor gamma. In addition, correlation between pre-stimulus beta and pre-response gamma in the motor cortex and post-stimulus frontal theta activity suggest bidirectional interactions between motor and frontal areas. These interactions likely underlie recruitment of preparatory and inhibitory neural activity during rapid motor control. Furthermore, pupillary and frontal theta effects during learned switches demonstrate that increases in inhibitory control of behavior can occur automatically, without conscious awareness.

**Significance Statement:** Goal-directed control is crucial for overriding habits and producing desired actions, which can fail during errors and accidents, and may be impaired in addiction, attention-deficit disorders, or dementia. This type of control, including response inhibition, is typically accompanied by frontal theta-band activity. We examined the relationship between frontal theta and response inhibition during unconscious pattern learning. First, we found that frontal activity was sensitive to changes in control and correlated with reaction times. Second, insufficient motor preparation predicted greater frontal activity, reflecting a greater need for control, which in turn predicted greater response-related motor activity. These results link the frontal and motor cortices, providing possible mechanisms for controlled behavior while demonstrating that goal-directed control can proceed automatically and unconsciously.

## 1. Introduction

Human behavior is often argued to be under both cognitive and automatic control. Cognitive control is thought to direct behavior through explicit knowledge and expectation. Investigations into the interplay of cognitive and automatic control often use speeded response tasks involving rapid streams of expected stimuli, where the intrusion of an unexpected stimulus would require cognitive activity to intervene in order to inhibit the automatic speeded response (Kramer et al. 2011). However, the distinction between “cognitive inhibition” and “automatic” responding has been questioned (McBride et al. 2012, Hommel 2013, Jasinska 2013), and may not involve altogether separate brain activity. Furthermore, although cognitive control and conscious awareness are thought to be tightly linked, motor control can be implicitly acquired and remain outside of conscious awareness (Kunde et al. 2012). Subjects and patients alike can implicitly learn rather complex motor sequences (Meissner et al. 2018), but automatic inhibition via unconscious learning has only recently been studied (Isabella et al. 2019). It remains to be determined whether automatic inhibition is supported by different patterns of brain activity than typically ascribed to cognitive control, or whether cognitive and automatic inhibitory control can be generated by similar neural processes.

It has been suggested that cognitive control originates in frontal cortex mediated by cortical oscillations that underlie long-range communication within the brain (Buzsaki et al. 2012, Cavanagh et al. 2014). A strong candidate for a physiological mechanism of cognitive control is frontal theta oscillations, which are known to increase during working memory (Jensen et al. 2002), mental arithmetic (Gartner et al. 2015), response preparation (Womelsdorf et al. 2010), response switching (Cheyne et al. 2012) and response inhibition (Isabella et al. 2015). In addition, frontal theta power was decreased and with shorter latency to peak on trials with faster, more automatic switch trials in a speeded response switching task (Cheyne et al. 2012). These findings suggest that theta activity may be indicative of the need for and timing of cognitive control. In addition, it has been suggested that the function of frontal theta is sensorimotor integration (Cruikshank et al. 2012). However this frontal activity may reflect a simple alarm signaling the need for cognitive control, without having a functional role in downstream signaling (Cavanagh et al. 2014). In order to achieve control of behavior, the brain must integrate cognitive control processes into sensorimotor systems. If frontal theta represents engagement of cognitive control, its relationship with sensorimotor cortex has not been established.

There is evidence that the sensorimotor cortex is sensitive to inhibitory signals and also to cognitive control processes. Increasing beta event-related desynchronization (ERD) has been linked with response selection (Donner et al. 2009), response certainty (Tzagarakis et al. 2010), response speed (Pastotter et al. 2012), and commission of response errors (Cheyne et al. 2012). Furthermore, previous work in our group demonstrated gamma event-related synchronization (ERS) was delayed during error trials over go trials, thereby disentangling this signal from motor parameters (Isabella et al. 2015). If there are signals within the sensorimotor cortex that coordinate with frontal theta in order to produce cognitive control over behavior, they are likely to include beta or gamma.

In the current study, in order to determine the exact relationship between frontal theta power and cognitive control, we manipulated requirements for cognitive control of motor responses using a task that induces motor pattern learning (Isabella et al. 2019), and quantified task demands for cognitive control by simultaneously monitoring magnetoencephalography (MEG) measures of oscillatory brain activity and task-evoked pupil responses (TEPR). Mean TEPR (mTEPR) is a measure with a well-established relationship to cognitive function (Kahneman et al. 1966), and latency to maximum TEPR (TMax) is related to the timing of response selection (Einhauser et al. 2010). We then compared signals within the frontal and motor cortices, as they related to cognitive control and behavioral output.

## 2. Methods

### Subjects

Sixteen healthy right-handed adults (8 females, range 22-31 years) participated in this experiment. All subjects were recruited from the Toronto area and provided informed consent using protocols approved by the Hospital for Sick Children Research Ethics Board. Subjects were compensated 60 CAD for their participation.

### Go/Switch Task

The Go/Switch task employed in this study was similar to that used in a previous study (Isabella et al. 2019). All subjects were presented with a rapid stream of digits from “1” to “4”, where each target had an equal 25% probability of occurrence. Each stimulus was displayed for a fixed duration of 0.4 seconds, followed by a stimulus mask (“#”) that was displayed for an additional 2 seconds until the presentation of the next digit, for a total inter-trial interval of 2.4 seconds (**Figure 1**). All the stimuli and the mask were isoluminant. The subjects were informed that they were performing a go-switch task, for which the default movement to stimuli 1, 2, or 4 was a button press with the right index finger, with instructions to switch response hands to the left index finger when presented with the target “3” stimulus.

**Figure 1.**
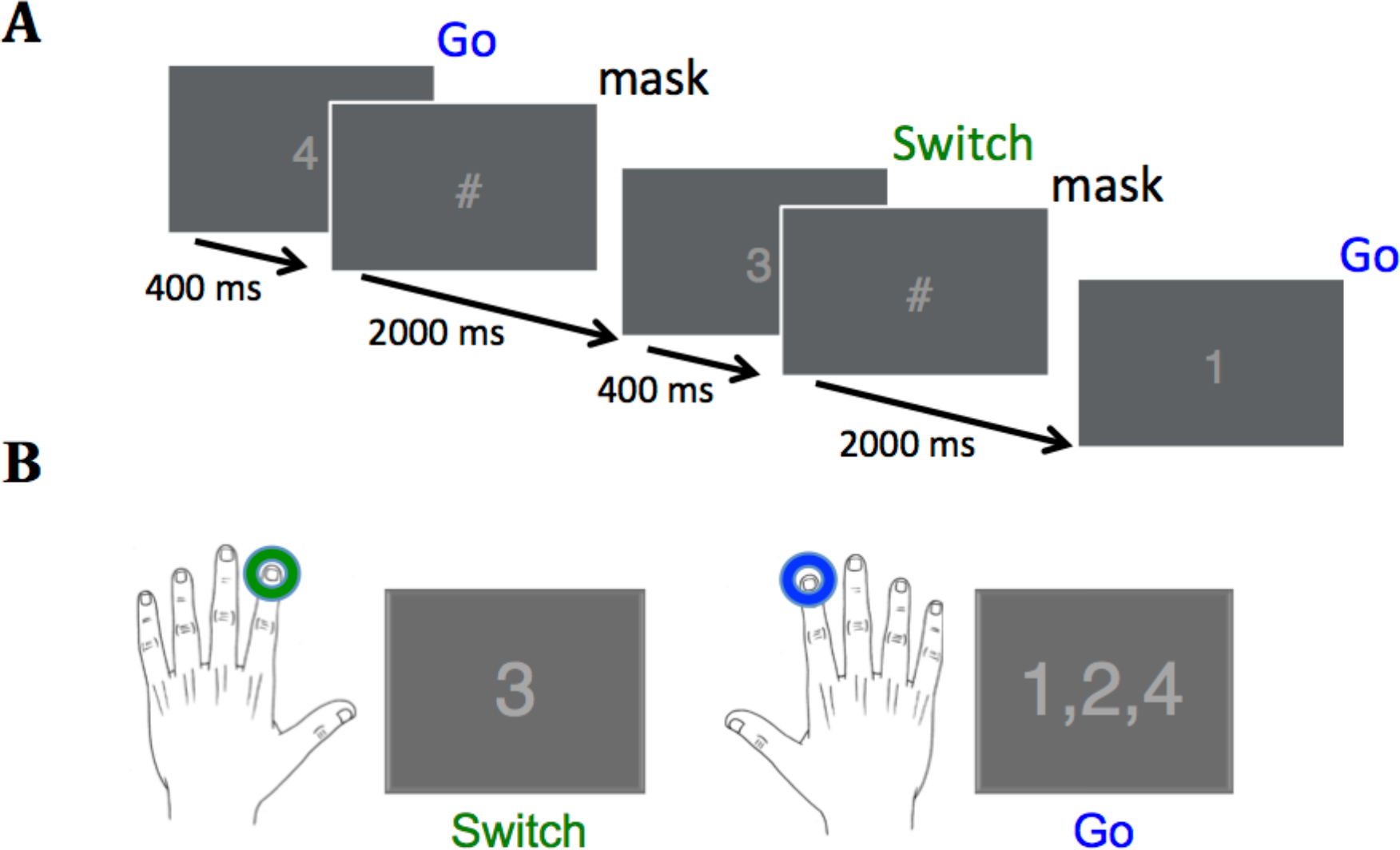
Task design and required responses. (A) Digits were presented every 2400 ms for a duration of 400 ms, followed by a stimulus mask (“#”) for a duration of 2000 ms. Overall probability for each of the four stimuli was 25%. (B) The correct response to the Switch stimulus (“3”) was a left index finger button press, whereas the correct response to all other stimuli was a right index finger button press. (reprinted with permission from Isabella et al., 2019).

Subjects performed this task over 244 trials across each of 6 blocks. Each block began with 4 trials containing stimuli (digits 1-4) chosen at random. Subjects were uninformed that the remaining 240 stimuli were presented in 30 repeats of an 8-trial probabilistic sequence (3-1-4-3-2-4-1-2), known to induce pattern learning in adults (Isabella et al. 2019) and also in typically developing children (aged 7–12 years) during a serial reaction time task (Gabriel et al. 2011). Stimuli for 90% of trials followed the sequence order (Pattern), whereas for the remaining 10% of trials, the stimulus for the individual trial within the 8-trial sequence did not follow the sequence order (Deviant).

Subjects were instructed to “respond as quickly as possible without committing too many errors”. They were informed that they would receive a $2 performance reward for each block with overall RT under 0.4 s and error rate under 30% for right (Go) and left (Switch) responses. Once the target RT and error rate were achieved, subjects were instructed to maintain or improve their scores in order to continue receiving the $2 monetary reward for each block (for a maximum reward of $12), in order to maintain motivation for the duration of the experiment. Before commencing the test blocks, subjects performed a practice block consisting of the same stimuli presented in random order, for 64 trials lasting 2.5 minutes. For the practice block only, feedback in the form of RT was presented on the left side of the screen after each response to help subjects determine their speed-accuracy strategy. Ahead of the test blocks, it was suggested to use the timing of the stimulus mask at 0.4 s as a benchmark for their RT on each trial.

Subjects were told that the goal of the study was to test the effect of a particular strategy, i.e., to covertly say the number as it appears on the screen. Following task performance on each of the 6 blocks, subjects were presented with the average reaction times and error rates for Go and Switch responses and were encouraged to improve their score for the next block. At the end of the experiment, subjects were asked for general feedback on the task, and to write out a sample stream of stimuli from the experiment. This was in order to assess whether the presence of the sequence of stimuli was explicitly learned.

### Recordings

Neuromagnetic activity was recorded using a whole head 151-channel CTF MEG system (MISL, Coquitlam, BC, Canada) in a magnetically shielded room. Data were collected at a rate of 600 samples/s. T1-weighted structural MR images were obtained from each subject using a Siemens 3T Magnetom Trio scanner. Small coils placed at fiducial locations (nasion and preauricular points) were used to monitor head position during recording and co-register source images to the subject’s MRI. Subjects sat upright in an adjustable chair and responses were collected using a nonmagnetic fiber optic response pad (LUMItouch Response System, Lightwave Medical Industries, Burnaby, Canada). Stimuli were presented using Presentation Software (version 14.9, www.neurobs.com) via a LCD projector on a back-projection screen.

Real-time TEPR was measured using an EyeLink 1000 system (SR Research, Ottawa, Canada), recording at 600 Hz and synchronized with the neuromagnetic activity. Pupil diameter was measured in arbitrary units.

### Analysis

#### i. Behavioral analysis

##### Response Types

Response types were defined as follows:

- Pattern Go (PGo): correct Go response (right index) to the Go stimulus (the digits 1, 2, or 4) matching the repeated pattern. I.e., occurring in the expected location within the sequence.
- Pattern Switch (PSw): correct switch response (left index) to the probable Switch stimulus (the digit 3) matching the repeated pattern.
- Deviant Go (DGo): correct go response (right index) to a Go stimulus (digit 1, 2, or 4) deviating from the repeated pattern, i.e., a Go stimulus where an expected “3” stimulus would have occurred requiring a Switch to a left index response. For example, given the intact pattern (3-1-4-3-2-4-1-2), a sample stream of stimuli with a Deviant Go trial (underlined) would be: 3-1-4-<u>1</u>-2-4-1-2. If the Pattern stimulus was expected, a change from a Switch to a Go response was required.
- Deviant Switch (DSw): correct switch response (left index) to improbable switch “3” stimulus, deviating from the repeated sequence, i.e., where the expected Pattern stimulus would have required a Go response. For example, a sample stream of stimuli with a Deviant Switch trial (underlined) would be: 3-1-4-3-2-<u>3</u>-1-2. For this response type, if the Pattern stimulus was expected, a change from a Go to a Switch response was required.

Importantly, all trial types as defined were preceded by a Go response. Any missed (nonresponse within 1.5 seconds of stimulus presentation) trials were rare and not included in any analyses. All trials following a “3” stimulus were not included in the analysis, as subjects quickly learned that the Switch stimulus “3” never occurred twice in succession and therefore could explicitly predict that a Go trial would follow a Switch trial. Any trials containing the incorrect response were also not included in the analysis.

##### Reaction Times

RT was measured as the difference in time between stimulus onset and the button press within 1.5 seconds of each trial. Any trials that did not include a physical button press or contained more than one button press were not included in the analyses.

##### Efficiency

Efficiency of each subject’s performance was determined by calculating the accuracy (percent correct) and dividing by the reaction time over the entire experiment for each of the 4 trial types (PGo, PSw, DGo, DSw).

#### ii. Pupil Diameter

Continuously recorded PD data was segmented into epochs and time-locked to stimulus onset. Eye blinks were linearly interpolated using a custom Matlab script, low pass filtered at 10 Hz, and then z-transformed within participants to minimize inter-subject variability, as performed in Smallwood et al, 2011 (Smallwood et al. 2011).

Pre-stimulus pupil diameter was measured as the mean z-scored pupil diameter for the 0.4 seconds preceding stimulus onset, which was then subtracted from the entire trial. mTEPR was measured as the mean z-scored pupil diameter for the 2.0 seconds following stimulus onset (until the subsequent pre-stimulus time period). In addition, the latency (in seconds) of maximum pupil dilation (TMax) was measured within the same time period following stimulus presentation.

#### iii. MEG analysis

Continuously recorded MEG data were segmented into epochs centered upon onset of the stimulus cue (cue-locked) and again centered upon the button response (response-locked) for each of the four response types described above. Localization of brain activity was carried out using frequency-based beamformer algorithms, using a single-sphere head model (Lalancette et al. 2011) implemented in the BrainWave Matlab toolbox developed at the Hospital for Sick Children (Jobst et al. 2018). Continuous head localization was used to monitor head motion throughout the recordings and trials were rejected off-line if head motion exceeded 5 mm.

We measured changes in induced cortical oscillations using the synthetic aperture magnetometry (SAM) algorithm (Robinson et al. 1999) in the frontal and sensorimotor cortices, in the theta, and beta and gamma frequency bands, respectively, using a priori regions and frequencies of interest pairings based on extant literature and previous findings (Cheyne et al. 2012, Isabella et al. 2015). Spatial normalization based on the MNI (T1) template brain was carried out using SPM8 (Wellcome Centre for Human Neuroimaging, London, UK). Talairach coordinates of peak activations were determined from the normalized images using the MNI to Talairach daemon (Lancaster et al. 2000). Group beamformer source images of the were superimposed onto a 3D rendered image of Colin-27 (CH2.nii) average brain using BrainWave and thresholded for demonstration purposes.

Whole-brain pseudo-t difference images were created by subtracting the source power during an active time window of 300 ms duration from a prestimulus baseline period of equal duration in the beta frequency band (15-30 Hz), 200 ms duration in the gamma requency band (60-90 Hz) or 500 ms duration in the theta (4-8 Hz) frequency bands. The size of these time windows were chosen in order to capture at least 2–4 oscillatory cycles of the lowest frequency of interest within a given window. The baseline time windows were set from 0.6 to 0.4 s preceding movement onset for the gamma frequency band, 0.8 to 0.5 s preceding movement onset for the beta frequency band, and 1.1 to 0.6 s preceding movement onset for the theta frequency band, in order to obtain a stable baseline for each. These baseline time windows were chosen based on empirical investigation of the data, in order to ensure the baseline begins after activity from the previous trial has terminated, and before activity in the current trial begins (Gross et al., 2013). This optimal baseline time period varied for different brain regions and frequency bands. The active time window was shifted in 50 ms increments from the period immediately following the baseline window, up to 0.5 s following the button-press response, and images were searched for the largest and most consistent peak activation within the region of interest in order to determine the source of the task-induced oscillatory activity within the frequency band of interest. That is, we sought to determine what were the greatest but stable task-induced effects for each frequency and location pairing.

To address our hypothesis of comparing task-induced source activity across trial types, time-frequency representations (TFRs) were constructed from source waveforms at the peak location determined above within the pseudo-timages. This was accomplished using a Morlet wavelet frequency transformation (Tallon-Baudary et al. 1997) of single trial source activity in 1 Hz steps using the following formula:

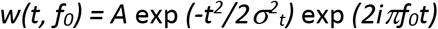

Wavelets are normalized so that their total energy is 1, using the normalization factor A that is equal to:

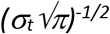

A convolution of the complex wavelet with the MEG signal is then derived and the magnitudec of this convolution used to create each TFR. This value is then converted to percent change in power relative to the same pre-movement baseline. Importantly, in order to exclude evoked activity, mean power was subtracted from the single trial power in the time-frequency plots, within-subject for each condition, to remove phase-locked activity and image primarily induced oscillatory activity (Keil et al. 2010). Mean power was then calculated for each subject over the time window of interest, for each trial type and TFR.

#### iv. Statistical analyses

RT was log-transformed to normalize its distribution. To examine differences between trial types across performance measures, TEPR, motor fields and mean oscillatory power, 2-by-2 within-subject repeated measures ANOVAs were conducted (factors were Switch and pattern), and any post-hoc comparisons were conducted using t-tests with Bonferroni corrections.

In order to investigate relationships between frontal theta and the other outcome measures of interest (TEPR, TMax, beta ERD, gamma ERS), a multiple regression approach was used in order to control for possible interactions of task variables (Sw and pattern). Relationships between measures were determined using the sum of squares from a repeated measures ANOVA between variables according to the following formula:

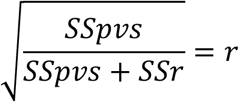

where SS*pvs* is the sum of squares for the predictor variables (as determined by the multiple regression), SS*r* is the sum of squares for the residual, and *r* is the correlation coefficient (Bland et al. 1995). All statistical tests were performed using R (Team 2017).

## 3. Results

### i. Behavioral Results

All 16 subjects complied with task instructions, completed all 6 blocks and provided feedback on the task. All subjects earned the maximum $12 performance reward. None of the subjects were able to replicate the complete stimulus sequence at the end of the experiment and failed to provide any evidence of explicit knowledge of the stimulus sequences.

#### Reaction Times

To determine the effects of task (Switch = Go/Sw and pattern = Pattern/Deviant) on responses, reaction times were measures as the duration between stimulus onset and the button press response for the four trial types of interest: PGo, DGo, PSw, and DSw. Mean RT was greater for Switch responses over Go, and greater for deviant over pattern trials (mean PGo = 0.347 s, DGo = 0.349 s, PSwitch = 0.370 s, DSwitch = 0.380 s, **Figure 2**). To determine the effects of task parameters on reaction times, a 2-way ANOVA was conducted on log-transformed averaged reaction times, revealing a statistical main effect of Switch (F(1,15) = 16.85, *p* < 0.0001) and of pattern (F(1,15) = 8.64, *p* = 0.01). post-hoc comparisons revealed significant differences between PGo and PSw, as well as PSw and DSw (all *p* < 0.003), but not PGo and DGo (*p* = 0.30). Switch responses of all types were delayed, but Deviant trials were delayed only for Sw trials and not Go trials (i.e. DGo). These results demonstrate an inverse relationship between response duration and variations in stimulus probability for all trial types except for the PGo-DGo difference.

**Figure 2.**
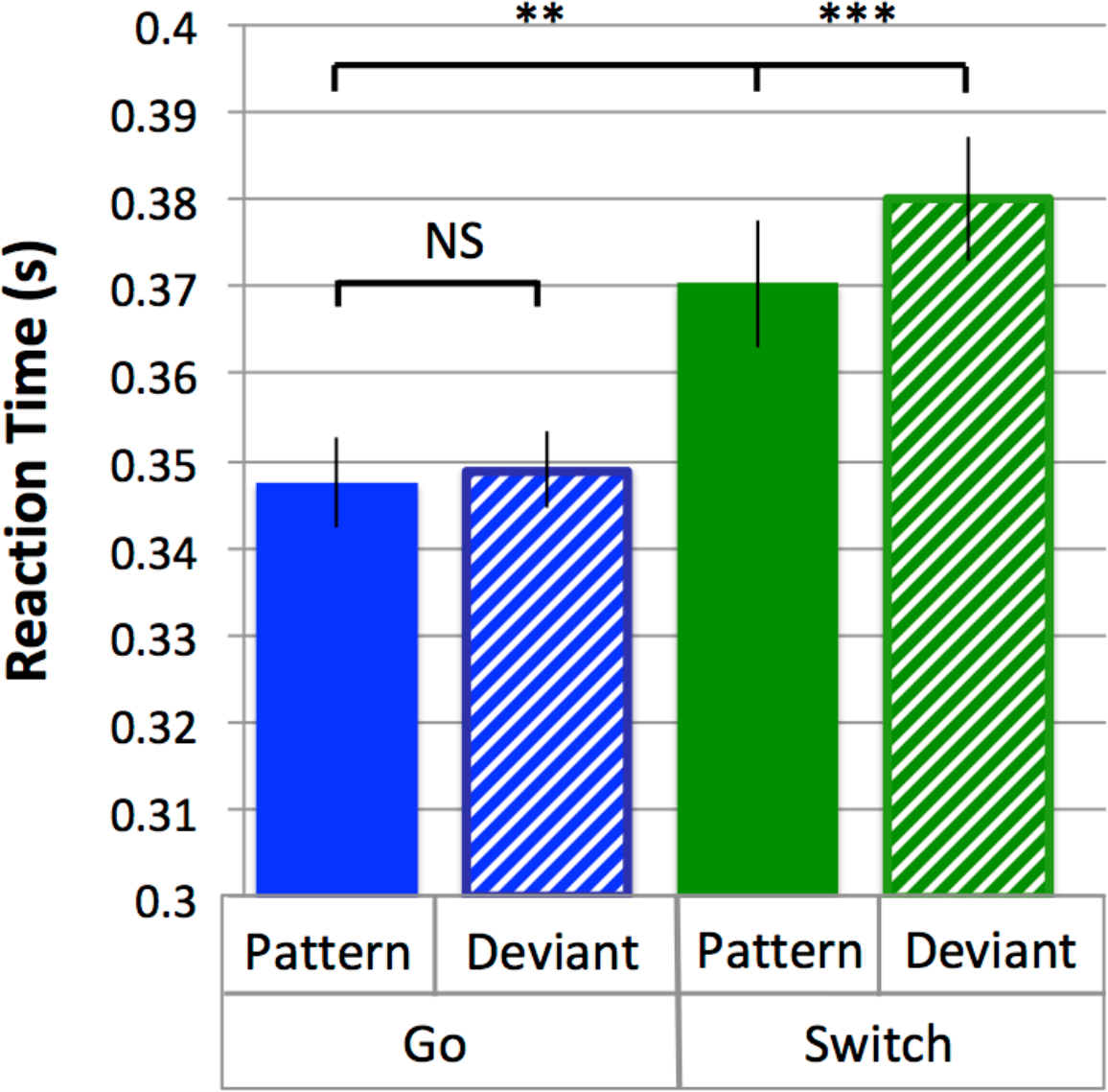
Reaction time. Mean RTs and standard errors for response types Pattern and Deviant Go, Pattern and Deviant Switch. Pattern Go, Pattern Switch and Deviant Switch response types were all significantly different from each other (all *p* < 0.003), but not Pattern and Deviant Go (*p* = 0.30).

#### Efficiency

Performance efficiency is defined as accuracy / reaction time, and reveals the overall speed-accuracy strategy utilized by each subject across trial types. Mean efficiency was greatest for PGo and DGo trials, and decreased for PSw and DSw trials (mean PGo = 2.83 correct/s, DGo = 2.85 correct/s, PSw = 2.31 correct/s, DSw = 2.15 correct/s; **Figure 3**). To determine the effects of task parameters on efficiency, a 2-way ANOVA was conducted on averaged efficiency rates, revealing a statistical main effect of Switch (F(1,15) = 23.35, *p* = 0.0002) but not of pattern (F(1,15) = 1.98, *p* = 0.18). Post-hoc comparisons revealed significant differences between PGo and PSw (*p* < 0.001), as well as PSw and DSw (*p* < 0.05), but not PGo and DGo (*p* = 0.30). Effects of task parameters on efficiency rates were similar to effects on RT, demonstrating that subjects did not change their speed-accuracy strategies across trial types. Subjects maintained consistent performance across P and DGo trials, with longer RT and efficiency for PSw and DSw, respectively.

**Figure 3.**
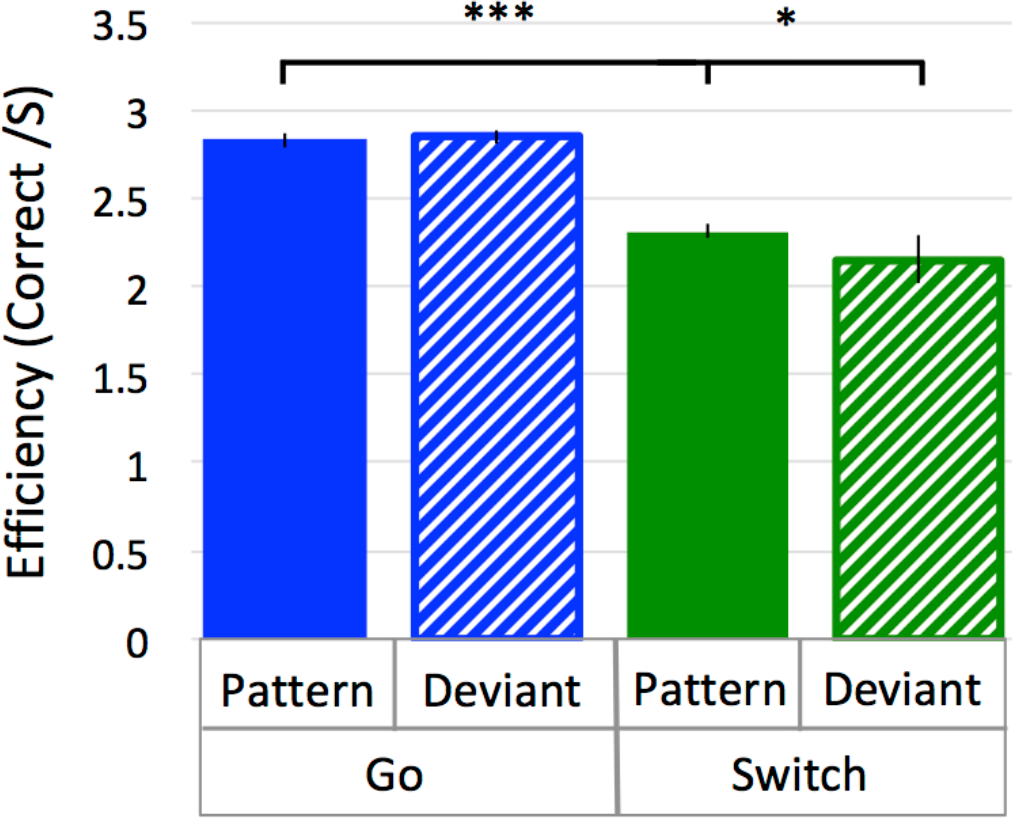
Efficiency Rates. Efficiency is the accuracy of each response divided by its RT, or, speed of correct responding. Mean efficiency and standard errors for all response types. Pattern Go, Pattern Switch and Deviant Switch response types were all significantly different from each other (all *p* < 0.05), but not Pattern and Deviant Go (*p* = 0.30).

### ii. Physiological Measures

#### Task-Evoked Pupil Responses

Pupil dilation is a well-established covert measure of quantifying cognitive control (Kahneman et al. 1966), and the latency to peak dilation (TMax) is also found to index relative latencies of response selection (Einhauser et al. 2010). In the current study, TEPR followed a typical time course, beginning at a minimum prior to stimulus onset, and dilating to a maximum diameter within 0.5 to 1.5 seconds (**Figure 4**). Diameters generally returned to approximately pre-stimulus levels following PGo trials ahead of the next trial at 2.4 seconds (n.b. all trials included in the analysis were preceded by Go trials). Mean TEPR was calculated as the mean baselined z-scored pupil diameter for 2 seconds following stimulus onset (time 0 to 2s), and was smallest for PGo trials, and increased for each of DGo, PSw and DSw trials (mean PGo = 0.25 ± 0.03 z, DGo = 0.43 ± 0.03 z, PSw = 0.46 ± 0.03 z, DSw = 0.57 ± 0.03 z; **Figure 5A**). In order to determine the effects of the task parameters on mTEPR, a 2-way ANOVA was conducted, revealing a statistical main effect of Switch (F(1,15) = 186.9, *p* < 0.001) and of pattern (F(1,15) = 88.72, *p* < 0.001). Post-hoc comparisons revealed significant differences between PGo and PSw, PGo and DGo, and Pattern and Deviant Sw (all *p* < 0.001). These results reveal a parametric increase in mTEPR with decreasing stimulus probability, and contrasted with RT results, consistent with previous findings that mTEPR and RT index different processes within cognitive control (Isabella et al. 2019).

**Figure 4.**
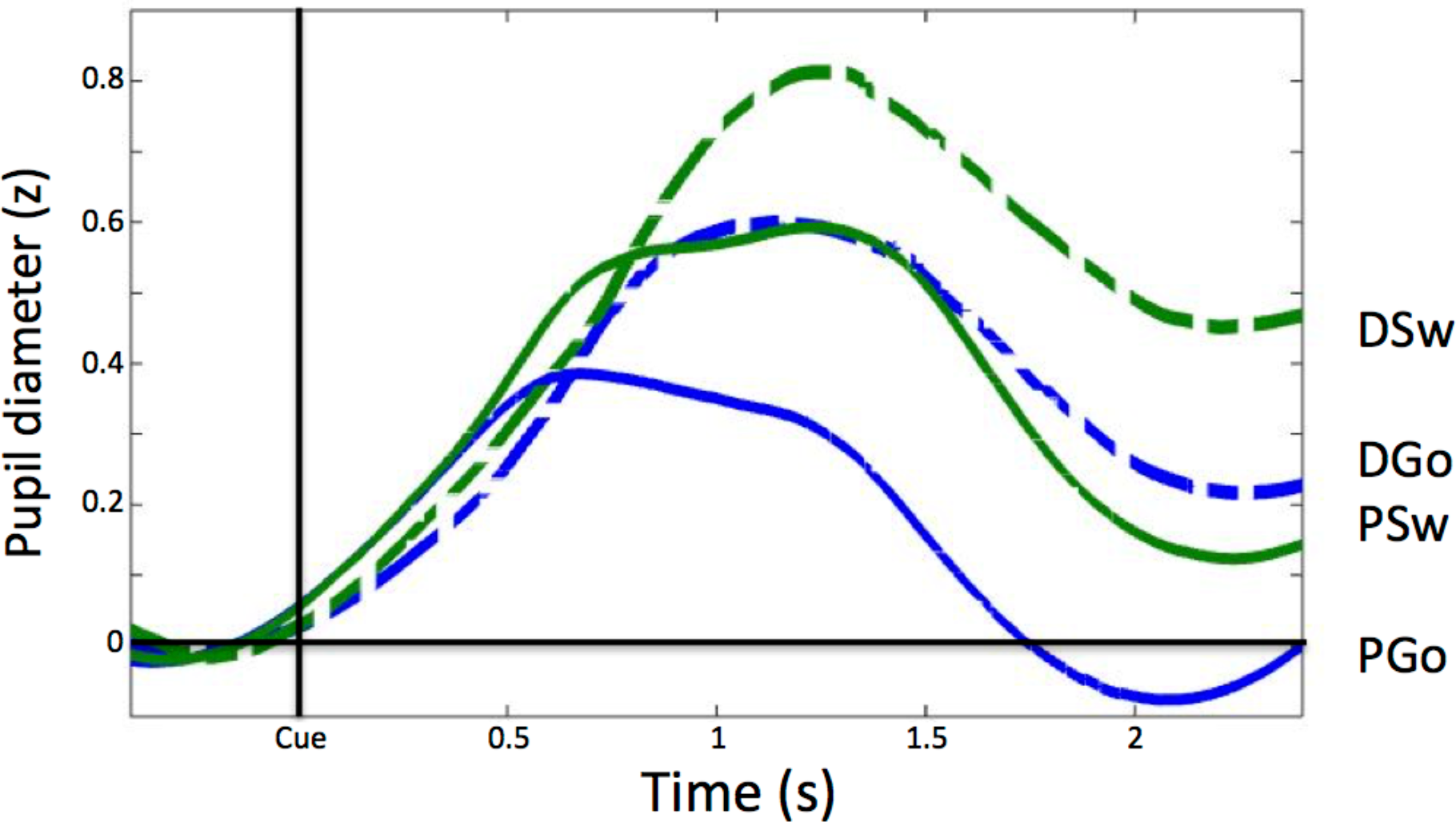
Pupil Diameter Time Course. Pupil diameter is within-subject z-scored, and timelocked to cue onset. Pre-stimulus PD for each trial was calculated as the average z-score value within the 0.4 seconds preceding stimulus onset, and was subtracted from the entire trial. Mean task-evoked pupil response for each trial type was calculated as the average z-score value over 2.0 seconds following stimulus presentation.

**Figure 5.**
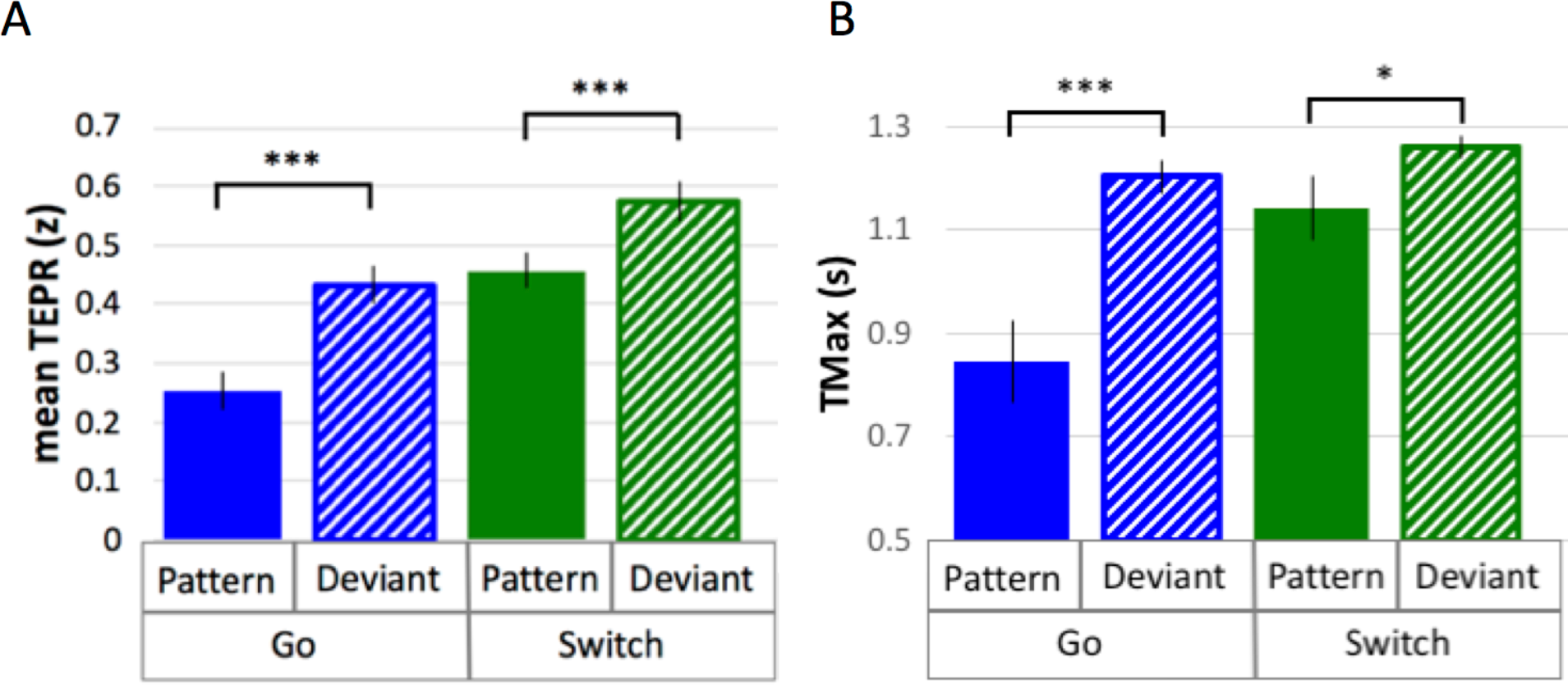
Task-Evoked Pupil Responses. **A.** Mean TEPR (in z-scores) and standard errors for all response types. All response types were significantly different (all *p* < 0.001) except for Deviant Go and Pattern Switch (*p* = 0.17). **B.** Mean latency to maximum pupil dilation (TMax; in seconds from cue onset) and standard errors for all response types. All response types were significantly different (all *p* ≤ 0.05) except for Deviant Go and Pattern Switch (*p* = 0.20).

In addition, TMax was calculated as the latency to maximum pupil dilation within the 2 seconds following stimulus onset, and similar to mTEPR, was fastest for PGo trials, and increased for each of PSw, DGo and DSw trials (mean PGo = 0.84 ± 0.08 s, DGo = 1.20 ± 0.03 s, PSw = 1.14 ± 0.06 s, DSw = 1.26 ± 0.02 s; **Figure 5B**). In order to determine the effects of the task parameters on mean TMax, a 2-way ANOVA was conducted, revealing a statistical main effect of Switch (F(1,15) = 36.36, *p* < 0.001) and of pattern (F(1,15) = 13.23, *p* = 0.002), with an interaction between the two (F(1,15) = 6.48, *p* = 0.02; **Figure 5B**). Post-hoc comparisons revealed significant differences between PGo and PSw (*p* < 0.0001), PGo and DGo (*p* < 0.0001), and P and DSw (*p* = 0.05). These results are in line with mTEPR findings, and most notable was the highly significant physiological differences in mTEPR and TMax between PGo and DGo trials, that were not observed in the behavioral responses RT and efficiency.

### iii. Neuromagnetic measures

#### Frontal Theta

The relationship between variations in cognitive control and frontal theta oscillations was of critical interest in the current study. SAM beamformer analysis revealed consistent theta band (4 – 8 Hz) oscillatory activity in the right middle frontal cortex (mean Talairach coordinates: x = 26, y = 59, z = 21, BA 10) for correct pattern and deviant trials, epoched to the button-press response (**Figure 6**; baseline = −1.1 to −0.6 s). To account for different trial numbers in each response type, PGo was used as a covariance dataset for all response types.

**Figure 6.**
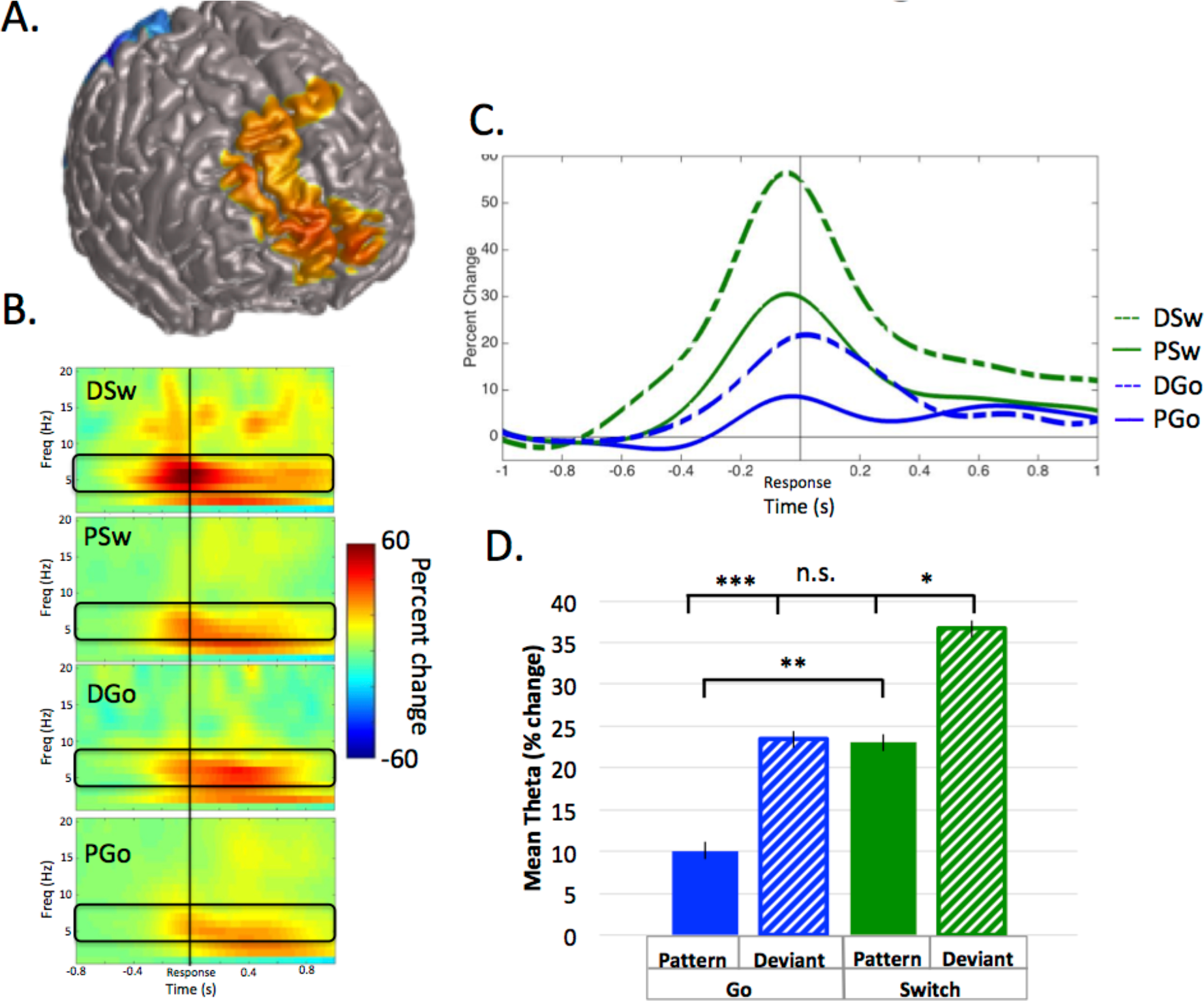
Frontal Theta Oscillatory Power. **A**. Source localization in right middle frontal cortex (x = 26, y = 59, z = 21, BA 10), with baseline set to −1.1 to −0.6 s relative to response onset. **B.** Time-frequency representations for all trial types, with the frequency bin of interest (4-8 Hz) outlined in black. C. Time courses for theta power in the contralateral motor cortex, for all trial types. D. Mean theta power (in percent change over baseline) and standard errors for all trial types. All response types were significantly different (all *p* < 0.019) except for Deviant Go and Pattern Switch (*p* = 0.21).

Theta power followed a typical time course, increasing to a maximum just prior to the response. Mean theta power was calculated as the mean percent change in power from 0.4 s prior to until 0.2 seconds after the button press response, relative to the pre-stimulus baseline. Theta power was smallest for PGo trials, and increased for each of PSw, DGo and DSw trials (mean PGo = 10.08 ± 2.16%, DGo = 23.39 ± 2.31%, PSw = 23.01 ± 2.35%, DSw = 36.64 ± 4.48%). In order to determine the effects of varied cognitive control on frontal theta oscillations, a 2-way ANOVA was conducted on mean power, revealing a statistical main effect of pattern (F(1,15) = 14.64, *p* = 0.002), and also a statistical effect of Switch (F(1,15) = 21.91, *p* < 0.001). Post-hoc comparisons revealed a difference between PGo and PSw (*p* = 0.002), between PGo and DGo (*p* < 0.001), and PSwitch and DSwitch, (*p* < 0.019). Interestingly, these results reveal theta has a similar relationship to task parameters as mTEPR and TMax, revealing a strong difference between PGo and DGo in the absence of any behavioural differences (RT or efficiency). This result supports our hypothesis that frontal theta is sensitive to parametric increases in cognitive control.

#### Sensorimotor Beta

Beta ERD is characterized by decreasing power preceding responses, but less so during uncertainty, and may be causally relevant to response execution. SAM beamformer analysis revealed consistent beta ERD oscillatory activity bilaterally in the sensorimotor cortex (mean Talairach coordinates left: x = −38, y = −21, z = 43; right: x = 38, y = −21, z = 40) for correct P and DGo and Sw trials, prior to responding (**Figure 7**; baseline = −0.7 to −0.4 s). We calculated the mean beta ERD power over two time periods: pre-stimmulus (−0.15 to 0 s) and pre-responses (−0.15 to 0 s). Mean power was calculated separately for each hemisphere.

**Figure 7.**
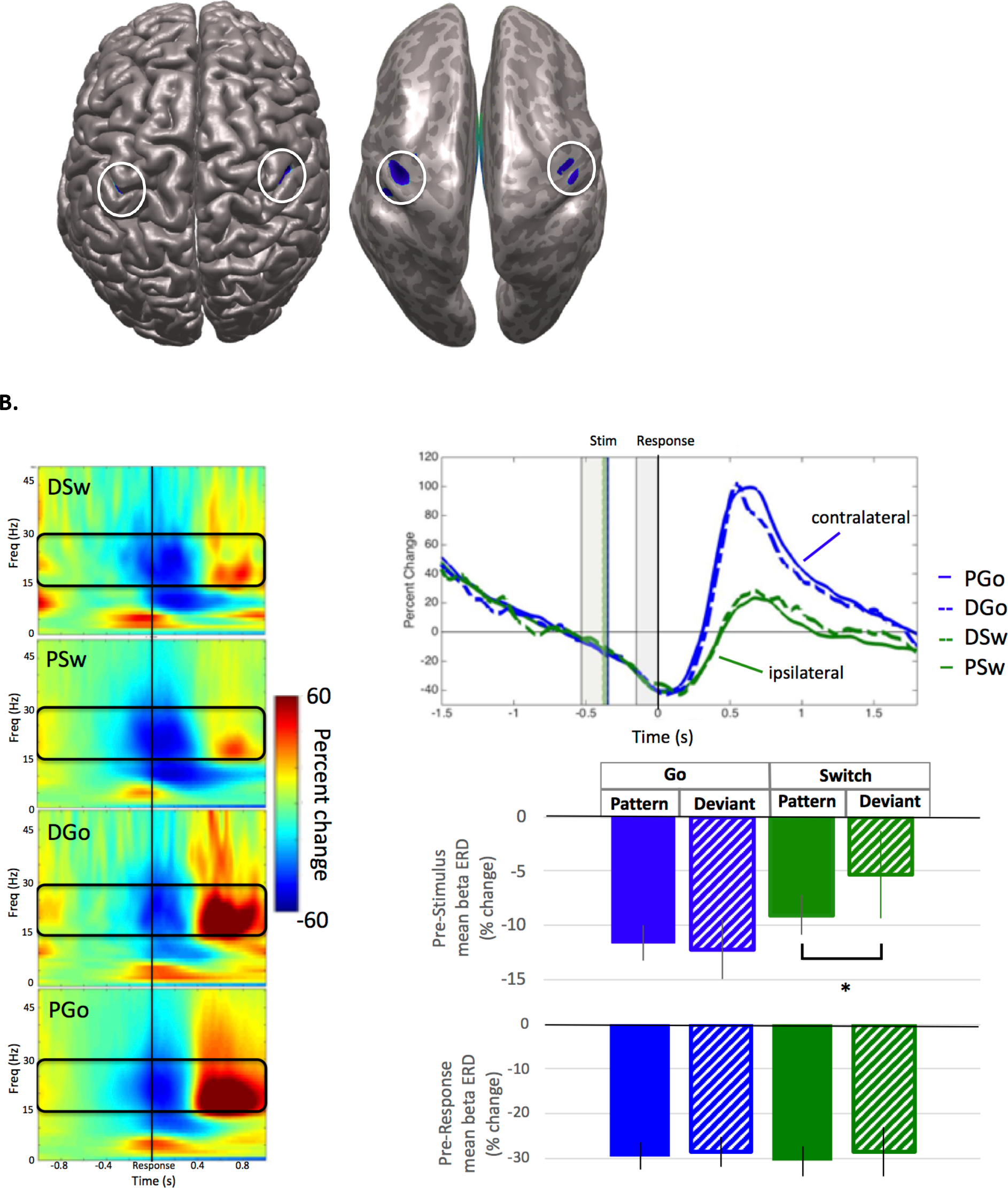
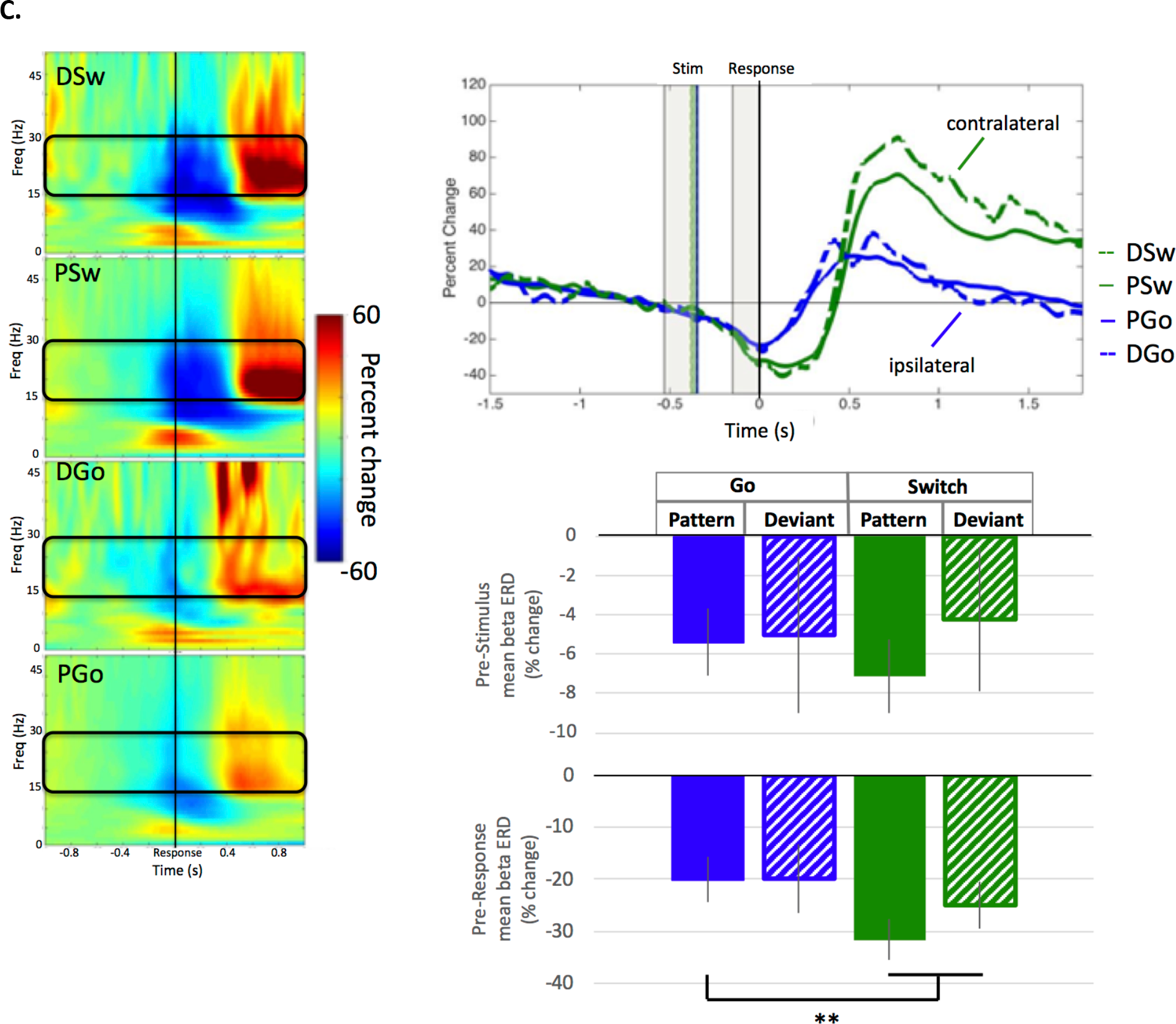
Pre-response beta event-related desynchronization. **A**. Source localization in left (x = −38, y = −21, z = 43; BA 3) and right (x = 38, y = −21, z = 40; BA 3) motor cortex, also shown on an inflated brain (right), with baseline set to −0.7 to −0.4 s relative to response onset. **B. Left motor cortex** (Left panel) Time-frequency representations for all trial types, with the frequency bin of interest (15-30 Hz) outlined in black. (Top panel) Time courses for beta ERD in the left motor cortex, for all trial types. (Bottom panel) Mean beta ERD (in percent change over baseline) and standard errors over 150 ms preceding response onset for all trial types. No significant differences were found. **C. Right motor cortex** (Left panel) Time-frequency representations for all trial types, with the frequency bin of interest (15-30 Hz) outlined in black. (Top panel) Time courses for beta ERD in the left motor cortex, for all trial types. (Bottom panel) Mean beta ERD (in percent change over baseline) and standard errors over 150 ms preceding response onset for all trial types. Post-hoc analysis revealed a significant increase for Pattern and Deviant Switch trials over Pattern Go trials (all *p* < 0.007), but not Deviant Go trials (*p* = 0.49).

Prior to stimulus onset, lateralization of beta ERD may reflect response preparation. In the left motor cortex, beta power decreased for all trial types (mean PGo = −11.6 ± 1.7%, DGo = −12.3 ± 2.8%, PSw = −9.1 ± 1.8%, DSw = −5.3 ± 4.0%, **Figure 7B** middle panel). In the right motor cortex, similar beta power decreases were observed for all trial types (mean PGo = −5.4 ± 1.7%, DGo = −5.1 ± 4.0%, PSw = −7.1 ± 1.8%, DSw = −4.2 ± 3.7%, **Figure 7C** middle panel). Given the high variability observed for the relatively low number of deviant trials, characterized by unstable within-subject virtual-sensor activity, deviant trials were excluded from further analysis. Paired t-tests revealed a significant difference in pre-stimulus beta ERD between PGo and PSw trials for the left (*p* = 0.024) but not the right motor cortex (*p* = 0.16), indicating reduced preparation for the right hand Go response ahead of anticipated PSw trials. The apparent overall effect of learning the stimulus pattern ahead of an anticipated Switch trial was not to increase Switch (right cortical) preparation as we had expected, but to reduce preparation of a Go (left cortical) response when it is less anticipated.

Prior to response onset, beta ERD in contralateral motor cortices may reflect delayed preparation for unexpected responses. In the left motor cortex, beta ERD was highly similar across trial types (mean DSw = −28.6 ± 5.4%, DGo = −28.6 ± 3.4, PGo = −29.62 ± 3.4%, PSw = −30.74 ± 3.47%, **Figure 7B** bottom panel). In the right motor cortex, beta ERD showed a sharp power decrease approximately 150 ms prior to responding, for Sw trials only (i.e. contralateral to response hand; mean PGo = −19.67 ± 3.76%, DGo = −19.9 ± 6.6%, PSw = −26.4 ± 5.2%, DSw = −24.9 ± 4.4%, **Figure 7C** bottom panel). In order to determine whether there were any effects of task parameters on pre-response beta activity, a 2-way ANOVA was conducted on mean power in the left motor cortex, with no significant effects of Switch (F(1,15) = 0.052, *p* = 0.8) or pattern(F(1,15) = 0.9, *p* = 0.3). Similarly, there were no significant effects in the right motor cortex (Switch F(1,15) = 3.44, *p* = 0.08; pattern F(1,15) = 0.09, *p* = 0.8). Inspection of the data revealed relatively high variability for DGo trials in the right motor cortex (ipsilateral), and may be driven by relatively low trial numbers (n=36/subject). Post-hoc analysis for the right motor cortex reveals differences between PGo and Sw responses (all *p* < 0.007), but not DGo (*p* = 0.49). These results demonstrate that left motor cortical oscillatory activity was similar across all trials types, but right motor cortical oscillatory activity was different between PGo and Sw trials. In combination with pre-stimulus beta ERD results, these results suggest that subjects reduced preparation of the prepotent Go response when a Switch trial is anticipated. This is in line with a model for attentional shift (Posner et al. 1990) where subjects must disengage from an attentional focus prior to shifting and engaging with a new one, in this case the Switch response. Furthermore, parallel activation of both responses may reflect a higher cost of inhibiting the prepared Go response over delayed activation of the Switch response.

#### Sensorimotor Gamma

Gamma ERS in the sensorimotor cortex has only recently been shown to vary with task parameters, and may be related to resolving response conflict. SAM beamformer analysis revealed consistent gamma ERS activity the sensorimotor cortex contralateral to the response hand (mean Talairach coordinates left: x = −34, y = −17, z = 43; right: x = 30, y = −10, z = 43) for correct P and D Go and Sw trials, commencing approximately 200 ms prior to responding (**Figure 8**; baseline = −0.6 to –0.4 s). In the ipsilateral motor cortex, there was no significant γ activity found (data not shown). We calculated the mean gamma ERS power in the contralateral motor cortex from onset until the response. Gamma ERS showed a similar pattern of effects as frontal theta, with the smallest ERS for PGo trials, and increasing for DGo, PSw, and DSw, respectively (mean PGo = 7.5 ± 1.8%, DGo = 13.7 ± 1.8, PSw = 25.1 ± 4.2%, DSw = −32.9 ± 4.5%). In order to determine the effects of task parameters on pre-response gamma activity, a 2-way ANOVA was conducted on mean power in the contralateral motor cortex, with significant effects of Switch (F(1,15) = 19.58, *p* < 0.001) and pattern(F(1,15) = 19.88, *p* < 0.001). Post-hoc comparisons revealed differences between PGo and PSw (*p* < 0.001), between PGo and DGo (*p* < 0.009), and PSw and DSw, (*p* = 0.003). This finding that, like frontal theta, sensorimotor gamma parametrically increases with decreasing stimulus probability is in line with our previous findings (Isabella et al. 2015), and suggests that sensorimotor gamma is sensitive to cognitive control. Given the similarity to frontal theta, sensorimotor gamma may be involved in integrating cognitive control signals into the motor cortex.

**Figure 8.**
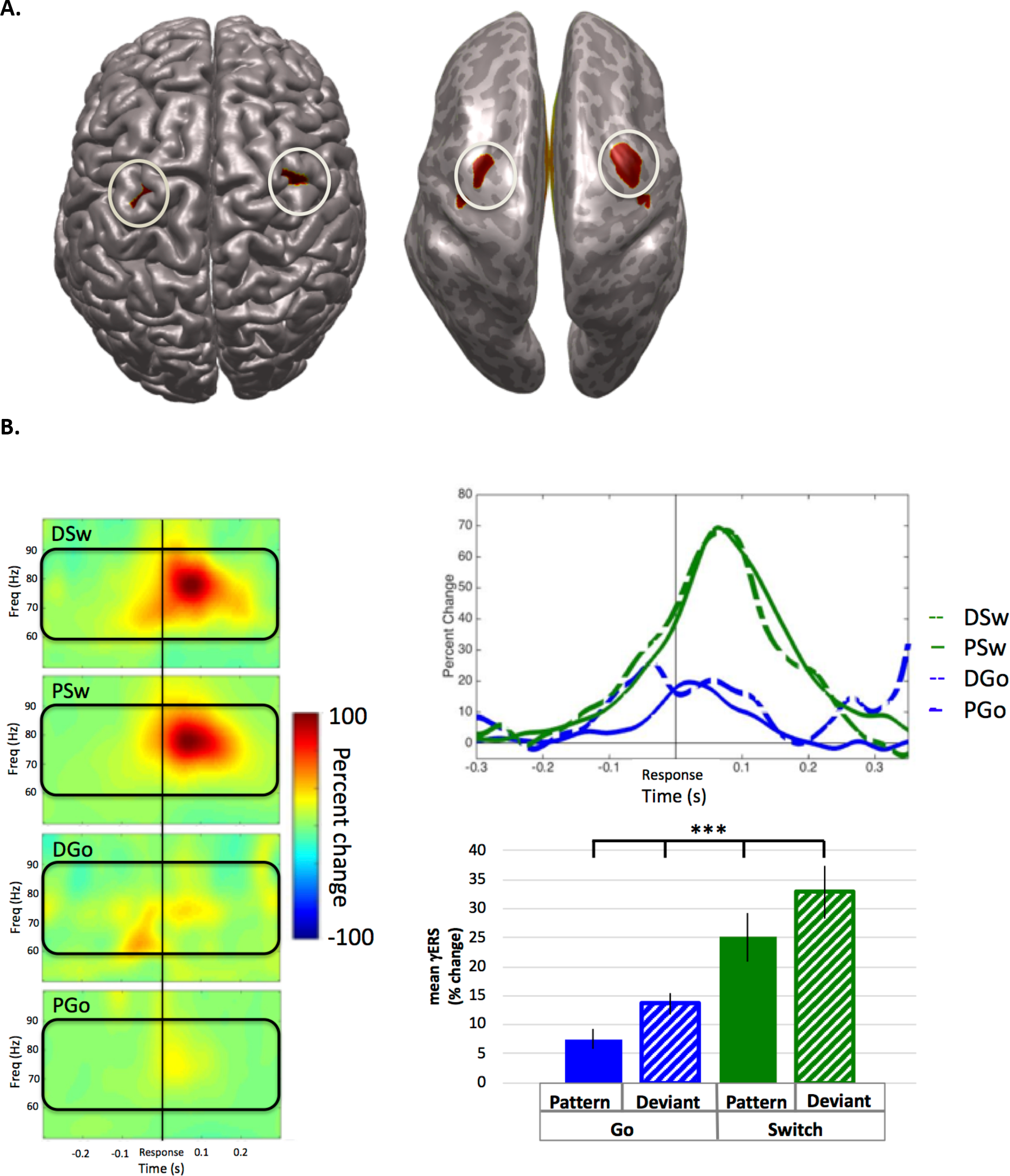
Pre-response gamma event-related synchronization. **A**. Source localization in left (x = −34, y = −17, z = 43; BA 4) and right (x = 30, y = −10, z = 43; BA 6) motor cortex, also shown on an inflated brain (right), with baseline set to −0.6 to −0.4 s relative to response onset. **B. Contralateral motor cortex** (Left panel) Time-frequency representations for all trial types, with the frequency bin of interest (60-90 Hz) outlined in black. (Top panel) Time courses for gamma ERS in the contralateral motor cortex, for all trial types. (Bottom panel) Mean gamma ERS (in percent change over baseline) and standard errors preceding response onset for all trial types. All response types were significantly different (all *p* < 0.003).

**Figure 9.**
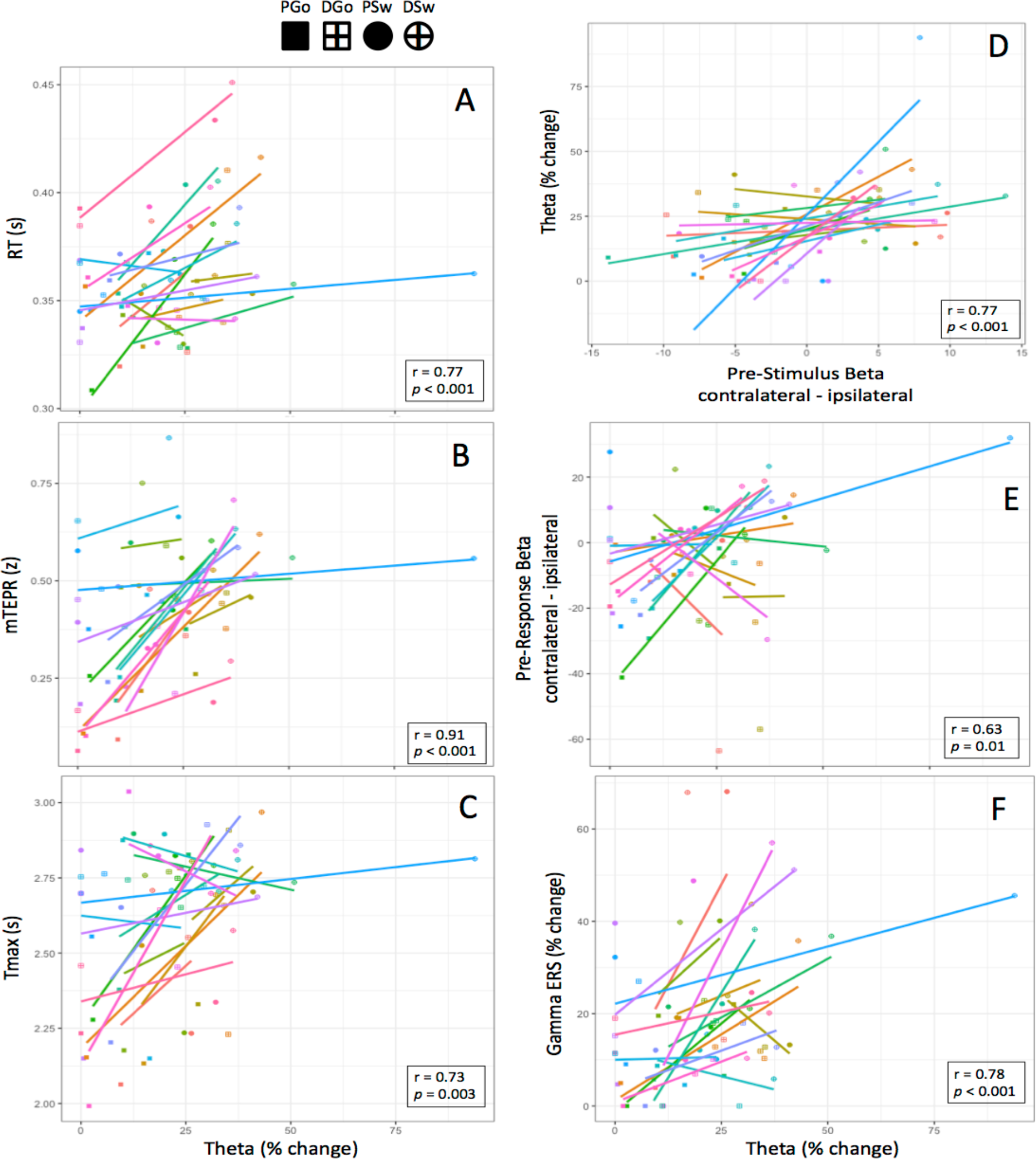
Regression Analysis. Anova tables were calculated within subject across repeated measures of Pattern and Switch. The sum of squares were analyzed to determine the strength (r) and significance (*p*) of covariance between frontal theta and the other outcome measures. **(A-F)** Data points are colour-coded for each subject, and symbols represent each trial type.

#### Regression Analyses

In order to investigate the relationship between frontal theta and the variety of measures in this study related to cognitive and motor control, we performed a multiple regression and analyzed the sum of squares to determine the strength and significance between the measures and within subjects. Results are shown in **Table 1** and **Figure 10**. Controlling for effects of pattern and Sw, there was a significant relationship between frontal theta and RT (r = 0.77, F(1,1,1,13) = 18.97, *p* < 0.001), mTEPR (r = 0.91, F(1,1,1,13) = 64.41, *p* < 0.001), pre-stimulus beta ERD (r_ipsi_ = 0.63, F_ipsi_(1,1,1,13) = 8.38, *p* = 0.013; r_contra_ = 0.56, F_contra_(1,1,1,13) = 5.88, *p* = 0.031), and pre-response gamma ERS (r = 0.78, F(1,1,1,13) = 19.71, *p* < 0.001). Additionally, there was a significant relationship between frontal theta and TMax, with an interaction between theta and pattern (r = 0.73, F(1,1,1,1,12) = 13.421, *p* = 0.003). There were no significant relationships between frontal theta and ipsilateral or contralateral pre-response beta ERD (**Table 1**).

**Table 1.** Anova tables. The ANOVA equation for each dependent variable was calculated, and the sum of squares for the predictors (SSpredictor) and residuals (SSr) are shown. These were used to calculate the r-values, F-statistic, and p-values for each regression.

| Dependent variable | ANOVA equation | SSpredictor | SSr | r | F | p |
| --- | --- | --- | --- | --- | --- | --- |
| RT | $RT \sim (\text{Theta} + \text{Go} + \text{P}) + \text{Error}(\text{subject}/(\text{Theta} + \text{Go} + \text{P}))$ | 0.005534 | 0.00379 | 0.770 | 18.97 | 0.0008 |
| mTEPR | $mTEPR \sim (\text{Theta} + \text{Go} + \text{P}) + \text{Error}(\text{subject}/(\text{Theta} + \text{Go} + \text{P}))$ | 0.30544 | 0.06165 | 0.912 | 64.41 | 0.0000 |
| Tmax | $TMax \sim (\text{Theta} * \text{P} + \text{Go}) + \text{Error}(\text{subject}/(\text{Theta} * \text{P} + \text{Go}))$ | 0.3432 | 0.30680 | 0.727 | 13.42 | 0.0033 |
| Pre-Stim Beta Ipsilateral | $\text{Theta} \sim (\text{Beta}_I + \text{Go} + \text{P}) + \text{Error}(\text{subject}/(\text{Beta}_I + \text{Go} + \text{P}))$ | 2235 | 3465 | 0.626 | 8.38 | 0.0125 |
| Pre-Stim Beta Contralateral | $\text{Theta} \sim (\text{Beta}_C + \text{Go} + \text{P}) + \text{Error}(\text{subject}/(\text{Beta}_C + \text{Go} + \text{P}))$ | 1234 | 2728 | 0.558 | 5.88 | 0.0306 |
| Pre-Stim Beta Contra-ipsi | $\text{Theta} \sim (\text{Beta}_{C-I} + \text{Go} + \text{P}) + \text{Error}(\text{subject}/(\text{Beta}_{C-I} + \text{Go} + \text{P}))$ | 3421 | 2313 | 0.772 | 19.23 | 0.0007 |
| Pre-Response Beta contralateral | $BERD_C \sim (\text{Theta} + \text{Go} + \text{P}) + \text{Error}(\text{subject}/(\text{Theta} + \text{Go} + \text{P}))$ | 162.2 | 1124.6 | 0.443 | 1.88 | 0.1940 |
| Pre-response Beta Contra-ipsi | $BERD_{C-I} \sim (\text{Theta} + \text{Go} + \text{P}) + \text{Error}(\text{subject}/(\text{Theta} + \text{Go} + \text{P}))$ | 2492 | 3723.0 | 0.655 | 8.70 | 0.0113 |
| Gamma ERS | $GErs \sim (\text{Theta} + \text{Go} + \text{P}) + \text{Error}(\text{subject}/(\text{Theta} + \text{Go} + \text{P}))$ | 2519.5 | 1662.1 | 0.776 | 19.71 | 0.0007 |

Although more challenging to interpret, given that analysis of lateralized readiness potentials is typically performed bilaterally (contralateral – ipsilateral), post-hoc analysis was performed on lateralized pre-stimulus and pre-response beta ERD in this manner. This analysis revealed a significant relationship between frontal theta and pre-stimulus lateralized beta ERD (r = 0.77, F(1,1,1,13) = 19.23, *p* < 0.001), as well as pre-response lateralized beta ERD (r = 0.63, F(1,1,1,13) = 8.70, *p* = 0.011). These analyses reveal a significant relationship between frontal theta and all of the other physiological and brain measures of interest. These widely distributed correlations with theta suggests a very meaningful role for this frontal signal within the cognitive control of behavior during this task.

## 4. Discussion

In the current study we manipulated cognitive control using a combined Go/Switch pattern learning task, without explicit awareness of the presence of a stimulus pattern. The task effects of Go/Switch and pattern/deviant led to increased RT, indicating pattern learning, and mTEPR confirming parametric variations in cognitive control. Increased cognitive control were associated with increased frontal theta, which was correlated with behavioural, pupillometric, and motor cortical activity. This provides the first evidence for a bidirectional relationship between frontal theta and the sensorimotor cortex, with important implications for the relationship between cognitive and motor control, discussed below.

### RT is insufficient for capturing cognitive control

Differences between RT and mTEPR are congruent with our previous findings (Isabella et al. 2019). Increased mTEPR has been associated with increased cognitive control (Kahneman et al. 1966), however, without a corresponding increase in RT or decrease in performance efficiency, we interpret this finding as increased cognitive effort without corresponding detectable behavioral effects. In particular, when comparing PGo and DGo, subjects increased cognitive effort to process the unexpected stimulus and response for DGo responses in the same amount of time as PGo responses. This finding has important implications for commonly used behavioral measures such as RT or efficiency in interpreting task difficulty or cognitive control, given that increasing cognitive effort need not produce detectable behavioral outcomes. We propose that mTEPR is a more sensitive and direct measure of cognitive control than RT or performance efficiency.

### Frontal theta is related to behavioral output via motor cortical signals

The current results include longer TMax during Sw and deviant than Go and pattern responses. Given that TMax has been shown to reflect the relative timing of response selection (Richer et al. 1983), in the absence of large differences in TMax, differences between Sw and deviant responses must lie within inhibitory and response preparatory processing. Preparation of Go responses was reduced for PSw trials, and thus pattern learning did not lead to greater or faster preparation of the Sw response, as indexed by beta ERD and TMax, respectively. Instead, pattern learning lead to reduced preparation and subsequently inhibition of the Go response. In contrast, increased cognitive effort to maintain consistent RT across the two types of Go responses was likely driven by frontal theta activity, which correlated with RT (inversely) and motor cortical activity. A relation between theta and RT is in agreement with a study showing that frontal theta power was decreased and onset delayed during stress, which correlated with longer RTs (Gartner et al. 2015). This suggests that speeded responses may depend on theta signaling, while being a finite resource. In addition, frontal theta power was proportional to working memory (Jensen et al. 2002) and also decreased on faster Sw trials (Cheyne et al. 2012), indicating that frontal theta increases are sensitive to the need for control processes. In the current study, frontal theta power was proportional to the amount of effort put into response inhibition and preparation, and this translated to behavioral output via a bidirectional interactions with the motor cortex.

We expected that theta would be related to the need for inhibition, as indexed by ipsilateral pre-stimulus beta ERD (**Figure 7**), and that theta would be inversely proportional to the extent of motor preparations, as indexed by contralateral pre-stimulus beta ERD, however the role for frontal theta was a function of both. Although beta ERD findings are not often reported as a contrast between the two hemispheres, these results show that cognitive control of motor output (pre-stimulus and pre-response) is a bilateral process. This is in line with recent findings that tDCS over left sensorimotor cortex reduces left and right no-go errors, highlighting the interrelations between bilateral sensorimotor cortices (Friedrich et al. 2018). Importantly, there is evidence for response conflict processing within the sensorimotor cortex (Coles et al. 1995, Cheyne et al. 2012), and it is hypothesized that this may be a role for gamma ERS contralateral to the executed movement.

Sensorimotor gamma ERS and frontal theta increased parametrically with cognitive control, and significant correlations were found between the two, suggesting that the gamma signal may be involved in integrating theta activity into the motor cortex. Furthermore, the information content of theta activity is likely related to updating the motor plan to the alternate response within the motor cortex prior to execution. This interpretation is congruent with our previous findings for sensorimotor gamma ERS when delayed gamma predicted error responses (Isabella et al. 2015). When competing responses are not sufficiently resolved in order to update the motor plan prior to responding, an error may occur. This interpretation is supported by evidence in clinical populations. Kurz et al. (2014) demonstrated increased sensorimotor beta ERD and decreased gamma ERS in children with cerebral palsy who had difficulty anticipating grip forces, possibly related to deficits in motor planning (Kurz et al. 2014). We speculate given the current findings that impaired motor planning may be related to deficits in signaling from the frontal cortex, or inefficient integration into the sensorimotor cortex via beta and gamma activity.

### Inhibitory control in the absence of awareness: role for frontal theta

Theta activity was localized to the right middle frontal cortex and increased from approximately stimulus onset, then peaked shortly prior to the response. As expected, theta power was sensitive to task parameters and distinguished trial types in a similar manner to mTEPR, albeit on a shorter time scale. Results of the regression analysis revealed a very strong correlation between these two outcome measures (r = 0.91). Previous studies have linked mTEPR with functional brain measures related to cognitive load and task difficulty, including alpha band power during a reading comprehension task (Scharinger et al. 2015) and theta power (evoked and oscillatory) during a combined flanker/n-back task (Scharinger et al. 2015). The current study identified a strong relation between mTEPR and oscillatory theta power and further demonstrated that these tracked parametric increases in cognitive control. Previous research has linked frontal theta with a variety of cognitive control processes, such as mental arithmetic (Gartner et al. 2015), response preparation (Womelsdorf et al. 2010), response switching (Cheyne et al. 2012) and response inhibition (Isabella et al. 2015). One study demonstrated the sensitivity of frontal theta to increasing load in a working memory task (Jensen et al. 2002). The current study supports the notion that frontal theta is sensitive to increasing load, and we extend those findings by demonstrating the relationship between theta and increasing cognitive effort using pupillometry.

Pupil diameter is tightly linked with activity in the locus coeruleus (LC) (Joshi et al. 2016), and noradrenergic and cholinergic pathways (Jepma et al. 2011, Reimer et al. 2016). Task-related pupil responses during cognitive processing were related to the BOLD signal and LC activity (Murphy et al. 2014, de Gee et al. 2017), and LC activity has been related to cognitive performance (Minzenberg et al. 2008). Therefore, it is likely that the current mTEPR findings are related to LC activity, and the tight relationship with frontal theta suggests the LC and frontal cortex are both engaged during focused task performance, while mTEPR could reflect subcortical interactions between the two.

Although top-down control of action is generally thought to occur within the cortex, it has been suggested that conscious and unconscious processes are implemented by the same neural substrates performing the same neural computations, and the difference between the two might only be a matter of degree (Horga et al. 2012). Others have gone further to suggest that there is no causal role for conscious processes in action control, and that automatic processes may underlie normal motor behavior that is generally attributed to top-down cognitive control (Kunde et al. 2012, McBride et al. 2012, Hommel 2013, Jasinska 2013). Current understanding of implicit learning suggests that it is also automatically acquired and not under cognitive control. That subjects in the current study were not consciously aware of the existence of a pattern suggests that they were automatically increasing effort required to inhibit responses, either when predicted as in PSwitch or when unpredicted as in DSwitch. The association of frontal theta power with inhibitory control, in the current study as well as in others, suggests that both inhibitory control and processing of other parameters associated with cognitive effort such as response selection may all be under automatic control.

## Conclusions

Frontal theta has been lauded as the ‘lingua franca’ for cognitive control (Cavanagh et al. 2014), however has lacked sufficient evidence to support this assertion, namely a link between behavioral and cognitive control. The current study provides a link between frontal theta and measures of cognitive control (mTEPR), as well as a bidirectional link with the sensorimotor cortex. This work provides evidence to support frontal theta as a mechanism for frontal control of behavior, via the motor cortex. In addition, that the stimulus pattern was learned and that subjects were not able to consciously repeat the stimulus pattern reveals that control of behavior occurred automatically via the same neural mechanisms, without conscious awareness.

## Acknowledgements

This work was supported by a Natural Sciences and Engineering Research Council of Canada Discovery Grant (#184018-09). We would like to thank Marc Lalancette for technical support.

